# Positive selection tends to act on exposed extracellular regions and delineate interaction modules targeted by pathogens

**DOI:** 10.64898/2026.09.21.753075

**Authors:** Laszlo Dobson, Éva Schád, Erzsébet Fichó, Ágnes Tantos, András Szabó, Gábor E. Tusnády, Rita Pancsa

## Abstract

**Background:** Positive selection shapes protein function by favoring amino acid substitutions that increase fitness. Although numerous studies have identified positively selected genes (PSGs), the structural principles underlying residue-level positive selection remain incompletely understood and previous studies have produced conflicting conclusions, particularly regarding the preferential location of positively selected residues (PSRs) within intrinsically disordered regions. In order to resolve these issues and to better understand the driving forces underlying positive selection in human proteins, we performed a comprehensive analysis on a manually curated, high quality collection of PSGs and PSRs as well as on a large-scale PSR dataset restricted to proteins with experimental structures.

**Results:** Human PSRs were significantly enriched in secreted and cell membrane proteins and preferentially localized to extracellular regions, solvent-exposed protein surfaces, coil structures, and protein-protein interaction interfaces, whereas neither domains nor intrinsically disordered regions showed an enrichment. Spatial clustering analysis reinforced that PSRs accumulate within localized surface patches. Integration of host-pathogen interaction data highlighted that adaptive changes preferentially affect extracellular molecular recognition surfaces, recurrently appearing in certain pathogen-interacting membrane protein families and particular domain types. Additionally, PSRs showed an enrichment in the residues of human-pathogen interaction interfaces that are in direct contact with pathogenic proteins, implying that neutralizing pathogen attacks is one of the driving forces behind the adaptive evolution of human proteins. By analysing human variation data we found that PSRs are enriched in benign but depleted in pathogenic substitutions, extending previous observations that positively selected genes contain elevated levels of missense variation. Notably, PSGs of the large-scale dataset were also enriched among clinical-stage drug targets, suggesting a potential link between positive selection and pharmacological relevance.

**Conclusion:** Together, our results provide a refined structural model of adaptive evolution in human proteins and identify extracellular exposed molecular recognition surfaces and human-pathogen interaction interfaces as recurrent hotspots of positive selection. Our results point to the direction that precise identification of human PSRs could delineate important host-pathogen interfaces that exert evolutionary selection pressure on humans and appoint novel targets with therapeutic potential.

## Background

During evolution, certain protein sites may undergo adaptive changes/positive selection that alter, or at least fine-tune function [1]. If the new function increases fitness, the relevant sites likely become subjects to purifying/negative selection. The functional specialization of redundant protein copies after gene duplication is one of the best described scenarios in which new functions can be sampled through adaptive evolution [2,3]. A plethora of studies focused on the identification of genes and functions subject to adaptive evolution in different organisms, and various approaches have been developed to detect them [4]. For instance, regions with significantly elevated substitution rates have been detected in the human lineage, called human accelerated regions [5–8], which turned out to be mainly non-coding genomic elements presumably of regulatory functions [6,8,9]. However, only a limited number of studies have focused specifically on protein-coding regions and how protein structure influences adaptive evolution [10–14] and most of them did not investigate human proteins but rather proteins of organisms with higher molecular adaptation rates [10]. Some related studies were performed in the early 2010s and therefore relied on less advanced methodology. They were also limited by the availability of experimentally determined protein structures, and because of this dependency, were inherently biased towards ordered domains (at the expense of intrinsically disordered regions (IDRs) [12,13]).

Among the early studies where the structural properties of positively selected residues (PSRs) were not (only) evaluated based on experimental structures, but sequence-based predictions to achieve a higher coverage of the proteome, a study claimed that in Drosophila species, residues within coils are more likely to be under positive selection pressure than residues in helices and beta-structures [15]. In plant protein families, PSRs were also claimed to be overrepresented in coil/turn residues and IDRs [16]. Another study proposed that positive selection preferentially affects IDRs in yeasts [17]. Some early studies were, however, burdened by misconceptions and methodological flaws, for instance they overlooked the fact that although IDRs lack a well-defined tertiary structure over time, they abundantly show propensities for secondary structure elements [18]. Thus they incorrectly handled disorder as a third secondary structure type over alpha helices and beta strands, basically confusing them with coils. Accordingly, in some of these studies dedicated disorder prediction methods were not used and their claims on the preference of PSRs for IDRs were inappropriately based on secondary structure predictions [15]. At the same time, a study of PSRs in Drosophila species has largely refuted the above results, since it proved that the identification of positively selected sites is not only highly dependent on the alignment method used, but is also subject to high false positive rates of over 50% coming from misaligned positions mistakenly inferred as being positively selected [19]. Since IDRs generally bear faster changes in evolution, they are particularly affected by misalignments [20–22], therefore a large fraction of the positions mistakenly flagged as PSRs likely belong to IDRs, which implies a strong bias towards disorder in the observed structural trends of PSRs [19]. More recently, in a dedicated study of *D. melanogaster* and *A. thaliana* population genomics data, Moutinho *et al.* showed that the impact of IDRs on adaptive mutation rates can differ between species [10]. Despite the caveats potentially corrupting the results of early studies and the substantial differences in the rates of molecular adaptation between different species [10], positive selection became widely attributed to disordered regions.

Finally, van der Lee *et al.* identified positively selected genes (PSGs) relevant to human biology by investigating protein-coding DNA sequences from nine simian (‘higher’) primates [23]. They applied a large-scale, robust comparative evolutionary analysis workflow to conservatively infer PSGs and codons in 11 170 high-quality one-to-one ortholog alignments, which successfully eliminated most issues affecting previous analyses. They were also the first to manually curate their data and exclude the PSRs that appeared to be artefacts (false positive PSRs) for several well-illustrated reasons, including misalignments [23]. The resulting dataset is of very high quality. The identified PSRs are remarkably enriched in proteins involved in innate and adaptive immunity, antimicrobial activity, reproduction, olfactory and taste receptors, which have long been known to be targets of positive selection in primates [6]. Also, transmembrane and secreted proteins were significantly overrepresented among PSGs, implying that recent positive selection acts on proteins that are in direct contact with environmental factors, such as pathogens and drugs [24]. In another study, Slodkowicz and Goldman identified PSRs from mammalian alignments and mapped those onto available protein structures [25]. For a subset of only twenty proteins with strong spatial clusterization of PSRs on the structures, the possible functional relevance of positive selection was evaluated in the context of the specific proteins [25].

In the latter two studies by van der Lee *et al.* and Slodkowicz and Goldman, the structural features of the identified PSRs (preferred localization in order/disorder, secondary structure, surface accessibility, low complexity etc.) have not been analysed. The interplay between adaptive evolution and the protein-protein interactions (PPIs) of human proteins is also of great interest and has also not been analysed by them in statistical terms. Since host-pathogen PPI interfaces are key targets of the evolutionary arms race between infectious agents and their hosts [26], proteins interacting with pathogens have repeatedly been shown to evolve under strong positive selection. In particular, viral interaction partners can account for a substantial fraction of adaptive evolution in mammals [27,28], but bacterial [28] and protozoan [29] pathogens have likewise been shown to exert strong selective pressure on the human genome. Since the two previously derived human PSR sets are of good quality and large enough, they provide an excellent opportunity to quantitatively assess the structural and functional principles underlying positive selection and how it affects PPIs (including interactions with pathogens) in human proteins.

In this work, relying on state-of-the-art computational methods including AlphaFold 2 [30] structure predictions, we performed a comprehensive analysis of the manually curated PSRs published by van der Lee *et al.* [23] and the structure-derived PSR set of Slodkowicz and Goldman [25]. These two datasets are expected to contain a relatively low fraction of false positive PSRs coming from misalignments due to the manual curation process filtering out erroneous PSRs and due to being restricted to protein regions with available structures, respectively. First, we explored the potential biases of the two datasets and then quantitatively evaluated the preferences of PSRs regarding protein localization, structural features (such as disorder, secondary structure and surface accessibility), as well as their relationship to pathogenic and benign human variations. Furthermore, we investigated whether the investigated PSRs preferentially affect human PPIs or human-pathogen interaction interfaces. Overall, our analyses complement previous observations and also revealed unexpected features of positively selected residues, some of which contradict previously reported trends. Based on our results, we also propose that the signal of adaptive changes could be considered when appointing novel therapeutic targets.

## Methods

In this section, first the two previously published PSR datasets are introduced that we re-use in this analysis, then the generation of respective random background sets is explained. We then list how the investigated characteristics, such as localization, structural, interaction and variation data, were annotated/obtained/predicted and reflected on the studied PSG/PSR protein datasets to perform comparisons against the respective backgrounds. Finally, we introduce the statistical tests used and the simple random forest classifier that we built to see if the identified features of PSRs distinguish them from other protein sites.

### Datasets

Positively Selected Residues (PSRs) were obtained from van der Lee *et al.* [23]. We verified all residues by mapping them onto the respective current UniProt sequences (UniProt [31] (2026_1 release)) and confirming that the mapped sequence positions matched the residue type reported in the original study. Mapping of the residues to canonical isoforms was preferred, but in the few cases when this was not possible, they were mapped onto alternative isoforms. 11 PSRs could not be mapped back to the reference proteome and were therefore excluded. The final curated dataset contains 922 residues in 325 proteins. We also used an additional dataset published by Slodkowicz and Goldman, where PSRs were identified from mammalian sequence alignments of protein regions with PDB structures using the criterion that PSRs should have statistically significant evidence of ω > 1 (where ω is the dN/dS ratio), resulting in 4,152 positions [25].

For both datasets we used an appropriate background set: for the high quality van der Lee set, of the 11 170 proteins from which the PSRs were derived 11 116 could be mapped to current UniProt sequences and these were used as the background pool (HQ_bg_pool). They reported that in this set some protein functions are strongly underrepresented (such as olfactory signaling and sensory perception of smell), while some others are moderately over– or underrepresented compared to the whole proteome. Therefore, by using this subset instead of the whole proteome for creating background sets (similarly to the van der Lee *et al.* analysis), we could avoid introducing biases potentially confounding the identified effects. For the Slodkowicz dataset PSRs were derived from proteins with PDB coverage (a pool of 3 348 proteins (LS_bg_pool). Residues were randomly sampled across all pool proteins while preserving both the protein-level and residue-level distributions observed in the PSR set. This procedure was repeated 10 times to generate the “random_background” sets. An analogous sampling strategy was applied to Positively Selected Genes (PSGs) to generate the “random_PSG” sets.

### Localization

Protein localization annotations were obtained from UniProt [31] and the Human Protein Atlas [32]. Additional evidence was incorporated from CCTOP [33] transmembrane topology predictions and the Surfy [34] surface protein predictor. Localization terms were grouped into six categories (nucleus, mitochondria, cytoplasm/cytosol, membrane, cell/plasma membrane, and secreted/vesicles). Proteins with any other localization were classified as unknown/other. For each protein, evidence from all sources was integrated using a scoring scheme. Because cytoplasmic annotations were common (many proteins localize to the cytoplasm during their life cycle), when the cytoplasm and another localization category had the same score the latter was favored. Similarly, membrane and cell membrane categories were resolved in favor of the cell membrane category, as cell membrane proteins are frequently annotated as membrane proteins. Predicted transmembrane segments or surface localization contributed additional support for membrane-associated classes. Proteins were assigned to the category with the highest overall score. In cases where no category or more than two categories shared the highest score, the protein was assigned an “unknown” localization. If two categories received the same highest score, a rule-based prioritization scheme was used. Membrane and cell membrane proteins lacking predicted transmembrane segments (according to CCTOP) were classified as Peripheral (cell) membrane proteins (see Supplementary Material 1: FigureS1).

### Structure

Structural and sequence-derived properties were assigned to all residues in the PSR and background datasets. PFAM [35] annotations were used to identify protein domains. Intrinsic disorder was predicted using AIUPred [36], while low-complexity regions were identified with SEG [37]. Linear motifs were retrieved from the ELM resource [38] and SLiMMine [39] predictions. Only high-confidence SLiMMine predictions (score > 0.9) with no annotation conflicts and at least one experimentally validated interaction partner were retained.

AlphaFold 2 (version 6) [40] protein structure models were used to determine structural properties. Secondary structure assignments and solvent accessibility values were calculated using DSSP [41]. Relative solvent accessibility (RSA) values were used to classify residues as exposed (RSA ≥ 0.36) or buried (RSA < 0.09) as described by Rost *et al.* [42].

Protein-protein interaction interfaces were identified using experimentally determined structures. Human protein residues were mapped to PDB [43] structures using SIFTS [44], and interchain atomic contacts were extracted from the 3did database [45]. Residues involved in at least one interchain contact in 3did were classified as contacting residues.

To assess the spatial clustering of residues, residue coordinates were partitioned using K-means clustering with k = 1, 2, or 3 clusters for proteins having at least 10 PSRs (this dataset includes 17 proteins and 305 positions from the HQ dataset). Cluster centroids were calculated from the residue coordinates and used as reference points. For each residue, the distance to the nearest centroid was determined and evaluated across a range of distance thresholds. To avoid considering weakly populated spatial regions, a centroid was considered valid only if at least W residues were located within the specified distance threshold from the centroid. Analyses were performed using minimum occupancy thresholds of W = 3, 4, and 5 residues.

### Interaction

Protein interaction data were obtained from IntAct [46], BioGrid [47], PDB [43] (using residue-level contacts from 3did [45]), HVIDB [48] and LeishMANIAdb [49]. Interactions were classified as either human-human or host-pathogen interactions. Pathogen interactions included viruses (taxid: 10239), bacteria (taxid: 2), and the following eukaryotic phyla containing relevant human pathogens: Fornicata (taxid: 207245), Parabasalia (taxid: 5719), Euglenozoa (taxid: 33682), Apicomplexa (taxid: 5794), Amoebozoa (taxid: 554915), Ascomycota (taxid: 4890), Basidiomycota (taxid: 5204), Mucoromycota (taxid: 1913637), Entomophthoromycota (taxid: 1264859)).

### Mutations

Missense mutations were obtained from UniProt variant resources (humsavar.txt and homo_sapiens_variation.txt.gz from https://ftp.uniprot.org/pub/databases/uniprot/current_release/knowledgebase/variants/). Only missense mutations with unambiguous annotations as either benign or pathogenic were retained for analysis.

To assess whether the amino acids introduced by the benign mutations affecting PSR positions appear in orthologs, we searched PSGs using a Quest For Orthologs approach (against the species used by van der Lee *et. al.* for generating the alignments from where PSRs were derived: *Pan troglodytes, Gorilla gorilla, Pongo abelii, Nomascus leucogenys, Macaca mulatta, Papio anubis* and *Chlorocebus sabaeus*) and aligned the sequences with ClustalO [50].

## Drug targets

To assess the pharmacological relevance of PSGs, human UniProt accessions were mapped to ChEMBL 37 targets [51], retaining only targets classified as SINGLE PROTEIN. Drug-target relationships were obtained from curated ChEMBL drug-mechanism annotations. Proteins linked to at least one compound in clinical development (with max_phase > 0) were classified as clinical-stage drug targets.

### Statistic tests

For each residue-level analysis, enrichment and depletion were assessed using contingency tables. Statistical significance was initially evaluated using Pearson’s χ² test. When any expected cell count was ≤5, Fisher’s exact test was used instead. Effect sizes were calculated as odds ratios and subsequently transformed to log₂ odds ratios for visualization and comparison. If any cell count was 0, 0.5 pseudocount was used to calculate odds ratios. For analyses involving randomized background datasets, statistical tests were performed independently for each randomization. To test the significance of the spatial clustering and solvent accessibility, two-sided Mann-Whitney tests were used at each threshold. For protein-level statistical analyses, such as protein subcellular localization and transmembrane classes, Pearson’s χ² tests were used in a similar manner, however the respective background pool dataset was used as control. Throughout the manuscript we consistently refer to enrichments as significant (p<0.05) or highly significant (p<0.01), but we deliberately do not burden the text by exact p-values, most of those can be found in figures, and all of them are available in the referenced supplementary tables.

### Predicting positively selected residues using random forest classification

To assess whether positively selected residues could be distinguished from non-selected residues based on their biological properties, a Random Forest machine learning approach was evaluated. To ensure strict separation between training and test data, protein-grouped cross-validation was employed for within-dataset evaluation, such that residues from the same protein were not shared between training and test sets. This procedure was repeated for each of the ten random-residue datasets for both HQ_PSR and LS_PSR. For cross-dataset evaluation, models trained on one dataset were tested on the other after excluding test proteins also present in the training set. The Random Forest model was trained using localization, membrane topology, solvent accessibility, secondary structure, interaction annotations, low-complexity regions, and predicted missense variant information as input features.

## Results

### Datasets

Positively Selected Residues (PSRs) were obtained from van der Lee *et al.* [23], with individual proteins potentially containing multiple PSRs. The van der Lee dataset is a high-quality, manually curated collection of 922 PSRs of 325 proteins (“HQ_PSR” set), that was derived from the subset of the human proteome where one-to-one orthologs could be unambiguously assigned and high-quality alignments prepared.

The 11 170 proteins whose one-to-one ortholog alignments were used to derive PSRs by van der Lee *et al.* were mapped to current UniProt sequences, resulting in a background pool of 11 116 proteins (“HQ_bg_pool” dataset). We prepared appropriate background sets by randomly sampling residues from this pool (“HQ_random_background” set), preserving both the protein-level and residue-level distributions observed in the PSR set; this sampling procedure was repeated 10 times. To gain an alternative background for specifically studying residue position-specific effects, we randomly selected protein residues from the set of Positively Selected Genes (PSGs, those genes that encode proteins with at least one PSR) themselves, preserving the residue-level distributions of the PSR set (“HQ_random_PSG” set), with 10 independent repetitions (Supplementary Material 2: Table S1). Because PSRs are unevenly distributed among PSGs and frequently accumulate within the same proteins, most analyses were performed at the residue level rather than the protein level. This approach captures PSR distribution across structural and functional categories and allows proteins with multiple PSRs to contribute proportionally to the analysis.

As an independent control, we also used the large-scale PSR dataset generated by Slodkowicz and Goldman [25], from which 4 129 PSR sites in 1 034 human proteins (“LS_PSR” set) could be mapped. Only 12 positively selected residues were common to both datasets, highlighting that PSR identification heavily depends on the phylogenetic scope of the underlying alignment, alignment method used and methodology of PSR detection. We generated random background sets in a similar manner as for the HQ dataset. The only exception is that for the Slodkowicz dataset the background pool contained those 3 348 protein regions (“LS_bg_pool” dataset) with experimental PDB structures that provided the basis of their PSR detection strategy [25]. Residues were randomly sampled from this pool to obtain the respective random background sets for the LS_PSR set (“LS_random_background” set). We also sampled random residues from the associated 1 034 PSGs as described for the manually curated set (“LS_random_PSG” set).

All analyses in the manuscript were performed on both datasets. Although it contains fewer PSRs, we consider the manually curated dataset from van der Lee *et al.* to be more reliable and free of obvious structural biases, and thus better suited for our analysis. Therefore, unless explicitly stated otherwise, the findings reported in the coming sections of the manuscript were derived from the van der Lee dataset (referred to as HQ_PSR dataset). The larger Slodkowicz PSR dataset (referred to as LS_PSR dataset) is limited to proteins with available PDB structures, thus it is inherently biased towards ordered protein segments. Also, it was generated through a fully automated pipeline that could potentially introduce a higher proportion of false positive or low-confidence PSR hits. Throughout the whole manuscript the two independent datasets are always compared to their respective backgrounds, the HQ_PSR dataset to the respective HQ_bg_pool and/or the two HQ_random datasets, and the LS_PSR dataset to the respective LS_bg_pool or the two LS_random datasets, therefore we will often just refer to “respective background(s)” for simplicity.

To assess the representativeness and potential biases of the datasets used to identify positively selected residues (PSRs), we calculated and compared several key characteristics of the background pool datasets and the human proteome (Supplementary Material 1: FigureS2). First, we found that the number of unique and total interaction partners is highly significantly elevated in the background pools. The LS dataset is focused on proteins with PDB structures, which may favor proteins having greater research attention, consequently they have been more extensively studied and have more documented protein–protein interactions. The enrichment in the HQ_bg_pool can be explained by its enrichment in proteins with developmental and neuronal functions, kinases and phosphatases, and proteins responsible for cell motility and migration. These proteins may have more extensive interaction networks than other proteins (Supplementary Material 3: Table S2). There is a highly significant increase in PDB coverage in the HQ_bg_pool compared to the human proteome, while the LS_bg_pool is fully covered by PDB structures (Supplementary Material 3: Table S3). In contrast, protein disorder results show an opposite trend, and the LS_bg_pool dataset has a highly significantly lower disorder content. The HQ_bg_pool is not biased towards proteins with lower disorder content compared to the whole proteome (Supplementary Material 3: Table S4), which shows that the criteria applied by van der Lee *et al.* for dataset assembly did not introduce considerable structural bias, and thus their PSR dataset is well-suited for studying the underlying structural determinants of positive selection in human proteins.

### Positively Selected Residues are enriched at the cell surface in bitopic membrane proteins

We developed an integrative pipeline that combines multiple data sources to assign a primary subcellular localization to each protein (see Methods, Supplementary Material 4: Table S5). We first examined the localization of proteins encoded by positively selected genes (PSGs). Consistent with the observations of van der Lee *et al.* [23], secreted and cell-membrane proteins were highly significantly enriched among PSGs compared to the respective background pool(HQ_bg_pool), while nuclear proteins were depleted (Fig. 1/A, Supplementary Material 5: Table S6-7). In contrast, Moutinho *et al.* reported elevated rates of adaptive evolution in nuclear proteins in *Drosophila melanogaster* and *Arabidopsis thaliana* [10], suggesting that not only the rate of adaptive evolution but also the preferential subcellular localizations of the affected proteins can differ among taxa.

**Figure 1.**
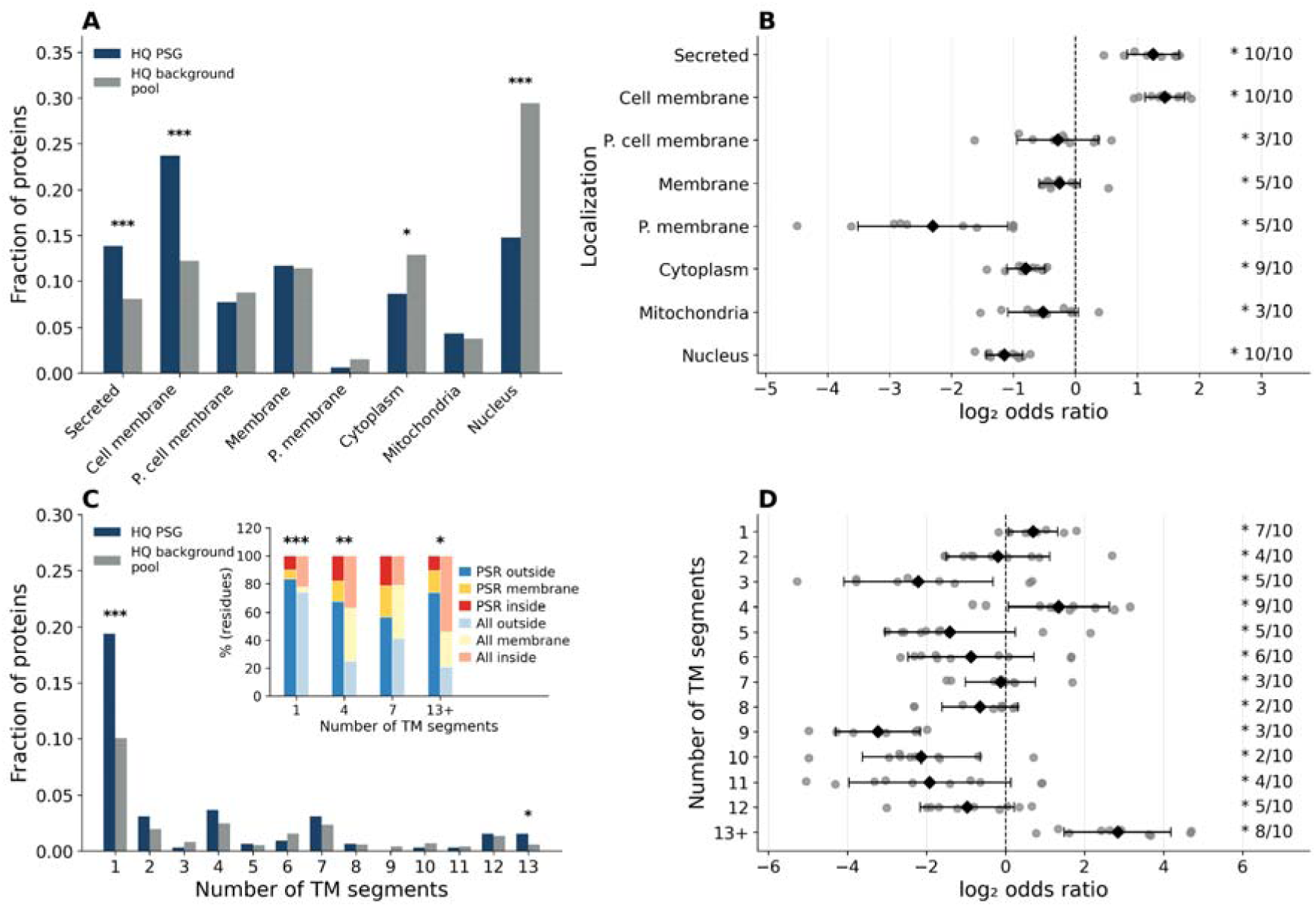
Positively selected residues are enriched in secreted and (mainly bitopic) cell membrane proteins. All figures show data for the HQ dataset. A: Distribution of protein localizations among positively selected genes (PSGs) compared to the HQ background pool proteins (P. membrane: Peripheral membrane). B: Forest plots showing the enrichment or depletion of PSRs in different localization categories compared to randomly selected background residue sets. C: Distribution of positively selected genes (PSGs) and HQ background pool proteins among different transmembrane topology classes. Inset: Distribution of topology labels (inside, membrane, or outside) for PSRs compared to all PSG residues in the topology classes with 1, 4, 7 and 13+ TMs. D: Forest plots showing the enrichment or depletion of PSRs in proteins belonging to different TM topology classes compared to randomly selected background residue sets. On the forest plots, individual dots represent enrichment estimates obtained from 10 independent randomization experiments. Diamonds indicate the mean log odds ratio (expected effect size), while error bars denote the standard deviation across randomizations. On the forest plots the accompanying text reports the fraction of randomizations yielding significant enrichment or depletion at p < 0.05 (*: significant) using Pearson’s χ² test. Positive log odds ratios indicate enrichment and negative values indicate depletion relative to the corresponding random background dataset. On the bar graphs * marks p < 0.05, ** marks p < 0.01 and *** marks p < 0.001 (Pearson’s χ² test).

At the residue level, we observed similar trends (Fig. 1/B). PSRs were enriched in secreted and cell-membrane proteins and depleted in nuclear and cytoplasmic proteins relative to the respective random background. These major localization patterns also appeared in the independent LS_PSR dataset, however with varying effect sizes (Supplementary Material 5: Table S8).

To test whether positive selection is associated with particular membrane protein topologies, we grouped transmembrane proteins (TMPs) by their predicted number of transmembrane (TM) segments. Among membrane PSGs, proteins containing a single TM segment (bitopic TMPs) represented the most abundant topology class, and PSGs showed a highly significant enrichment in bitopic TMPs compared to the respective background pool (Fig. 1/C, Supplementary Material 6: Table S9-10). At the residue level, PSRs were strongly enriched in bitopic TMPs (Fig. 1/D, Supplementary Material 6: Table S11), which became particularly pronounced when the analysis was restricted to cell-membrane proteins (Supplementary Material 6: Table S12). Additionally, PSRs were enriched in TMPs containing 4 and 13+ TM segments (Fig. 1/D). These patterns were not completely reproduced in the PDB-based LS_PSR dataset (maybe due to the general underrepresentation of TMPs in the PDB [52]), where PSRs were enriched in bitopic TMPs but depleted in TMPs with 4 and 13+ TM segments relative to the structure-mapped background. Other topology classes contained relatively few PSRs, limiting the power to detect consistent differences.

We next examined the position of individual PSRs relative to the membrane. In the topology classes where an enrichment of PSRs could be detected (those with 1, 4 and 13+ TM segments), PSRs are preferentially located on the extracellular (“outside”) side of the membrane (after normalization with the number of residues inside, in TM segments and outside) (Fig. 1/C, inset, Supplementary Material 6: Table S13-14).

Thus, positive selection pronouncedly affects the extracellular segments of bitopic TMPs, and TMPs with 4 and 13+ TM segments. TMPs with 4 and 13+ TM segments mainly come from the membrane-spanning 4A MS4A/tetraspan-related families and solute carriers (SLCs)/voltage-gated ion channel (VGIC) families, respectively, where the fraction of extracellular residues is <30%, but the fraction of outside PSRs is >60%. Members of the MS4A family are selectively expressed in immunocompetent cells and modulate the activity of immune receptors, such as pattern recognition receptors (PRRs) and Ig receptors. As such, they play a key role in different pathological settings, such as infectious diseases and cancer, which provides an explanation for their enrichment in PSRs compared to other TMP classes [53]. SLCs and VGICs are probably affected by positive selection because some of them act as obligate viral or bacterial receptors (e.g., SLC1A5 for retroviruses [54,55], or various SLCs used for nutrient scavenging by intracellular bacteria [56]) and/or because many of them are involved in the transport of nutrients, ions, metabolites, toxins and environmental xenobiotics, and in changing environments/during dietary shifts, environmental adaptation directly shapes transport physiology. By contrast, in 7TM proteins, which are predominantly G-protein-coupled receptors (GPCRs), the extracellular enrichment of PSRs was not significant in the HQ_PSR dataset (Fig. 1/C inset), but was significant in the LS_PSR dataset (being important drug targets [57], GPCRs are relatively well-represented in the PDB compared to other TMP families). A non-negligible fraction of PSRs localizing to their transmembrane segments could imply that the activation-dependent coordinated structural rearrangements within the TM regions of some GPCRs might be subject to recent adaptive evolution. Together, these results indicate that positive selection tends to preferentially affect secreted proteins and the extracellular regions of TMPs.

### Positively selected sites cluster on exposed surfaces

To investigate the structural determinants of positive selection, we compared sequence– and structure-derived properties of positively selected residues (PSRs) with those of randomly sampled background residues. It is important to note that since the Slodkowicz LS_PSR dataset was derived from PDB structures, it is expected to be inherently biased towards domains and ordered protein segments with very low representation of IDRs and especially low-complexity regions. In the high-confidence dataset, PSRs showed no consistent enrichment in PFAM domains or intrinsically disordered regions (IDRs) and were depleted in predicted low-complexity regions when compared with either respective backgrounds (HQ_PSR vs HQ_random_background and HQ_random_PSG). In the large-scale dataset (LS_PSR), the results seemed inconclusive: PSRs were depleted in PFAM domains relative to the structure-mapped residue background but enriched relative to residues sampled from PSGs (Supplementary Material 7: Table S15). These observations contrast with earlier studies, where PSRs were reported to be enriched in predicted IDRs in some species [17], although due to the substantial differences in the rates of adaptive evolution between those species and primates/human, matching structural preferences of PSRs are not necessarily expected.

Although PSRs showed no overall enrichment in PFAM domains, several domain types/families were recurrently found to contain PSRs in multiple proteins. The most frequently represented families were Sushi repeat domains (27 PSRs in 5 proteins), Trypsin domains (17 PSRs in 6 proteins), Immunoglobulin domains (13 PSRs in 7 proteins), and C-type lectin domains (11 PSRs in 4 proteins). Several of these families were also frequently represented in the LS_PSR dataset (Supplementary Material 7: Table S16). These domain families are predominantly found in secreted and cell-surface proteins, consistent with the observed enrichment of PSRs in extracellular proteins. Thus, while positive selection does not preferentially target protein domains in general, it repeatedly affects specific extracellular domain families mediating interactions with the external environment.

AlphaFold-predicted structural models allowed us to perform a comprehensive structural analysis of all PSRs despite the lack of available PDB structures (Supplementary Material 8: Table S17). At the level of secondary-structures, PSRs tend to be highly significantly depleted in α-helices and significantly enriched in coil regions in both datasets, whereas they showed slight enrichment in β-structures in the HQ_PSR dataset and slight depletion in the LS_PSR dataset compared with the respective background pools (Fig. 2/A,B, Supplementary Material 8: Table S18). PSRs were also highly significantly enriched among solvent-exposed residues (Fig. 2/C, Supplementary Material 8: Table S19), that agrees with previous observations in *Drosophila melanogaster* and *Arabidopsis thaliana*, where Moutinho and colleagues reported that adaptive substitutions preferentially occur on accessible protein surfaces [10]. The remarkably larger fraction of residues with high relative solvent accessibility in the HQ_PSR dataset compared with the structure-based LS_PSR dataset well reflects the unbiased nature of the former, wherein high-accessibility IDRs are also fairly represented (Fig. 2/C). Furthermore, after mapping PSRs onto PDB structures of complexes with multiple protein chains, we found that PSRs are significantly enriched in contacting residues of direct protein-protein interactions. Overall, human PSRs were found to be significantly enriched in exposed, interacting and coil residues and highly significantly depleted in α-helical residues compared with both respective backgrounds in both datasets (Fig. 2/D,E, Supplementary Material 8: Table S20). Interestingly, in the LS_PSR dataset, we did not observe an enrichment in exposed residues when comparing PSRs with the residue sets sampled from PSGs (LS_PSR vs LS_random_PSG), suggesting that this feature is might at least partly influenced by the selection of proteins represented in the dataset. Although AlphaFold2 predictions may be biased by the inclusion of experimental structures in its training data or template library, we observed the same structural characteristics after removing potentially affected structures [58] (Figure S3, Supplementary Material 8: Table S21).

**Figure 2.**
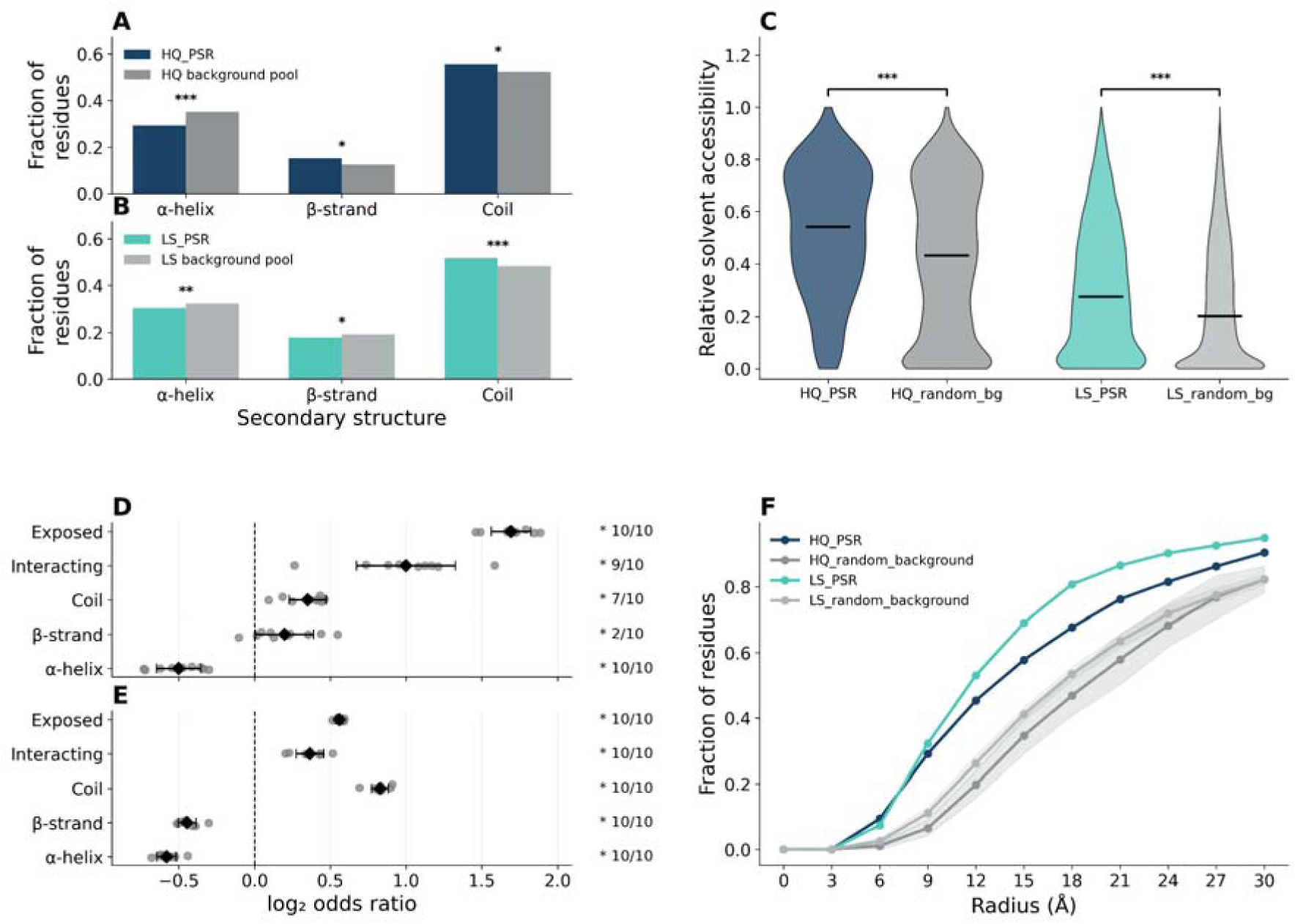
PSRs are enriched in coil structures, accessible surfaces and contacting residues of PPI interfaces. A and B: Comparison of assigned secondary structure types between positively selected residues (PSRs) and residues of the respective background pools for A: the HQ_PSR dataset and B: the LS_PSR dataset. C: Distributions of relative solvent accessibility values for the PSRs and respective random background residues for both datasets (random_bg: random_background). D and E: Forest plots showing the enrichment or depletion of secondary structure types, solvent-exposed residues, and contacting protein-protein interface residues among PSRs compared with the respective randomly sampled background residues in D: the HQ_PSR dataset and E: the LS_PSR dataset. For a description of forest plots see Figure 1. F: Fraction of residues assigned to hotspot regions as a function of the clustering radius (W = 3, k = 3) for PSRs and respective randomly sampled background residues for both datasets. Shaded areas indicate the standard deviation across randomization experiments. On the bar graphs (Pearson’s χ² test) and violin plots (two-sided Mann-Whitney tests) * marks p < 0.05, ** marks p < 0.01 and *** marks p < 0.001 significance levels.

To investigate whether PSRs form spatial clusters on protein structures, residue coordinates were analyzed using K-means clustering with one to three cluster centers. We analyzed proteins containing at least 10 PSRs, corresponding to 17 proteins in the HQ_PSR dataset and 105 proteins in the LS_PSR dataset (Supplementary Material 8: Table S22). Consistent with the findings of Slodkowicz and Goldman [25], in both datasets, PSRs formed spatial clusters more frequently than the residues in the respective randomized backgrounds. For example, using three cluster centers, a distance threshold of 12 Å, and a minimum of three residues per cluster, 45.4% of PSRs in the HQ_PSR dataset and 53.1% in the LS_PSR dataset were associated with clustered regions, approximately twice the fractions observed in the corresponding randomized backgrounds. Similar patterns were observed across a broad range of clustering parameters (Supplementary Material 9: Table S23, Fig. 2/F). Thus, despite substantial differences in the composition and ascertainment of the two datasets, PSRs consistently formed localized structural hotspots, suggesting that spatial clustering is a general feature of positively selected residues most likely bearing functional relevance.

### PSRs are enriched among residues directly contacted by pathogens

To investigate whether positive selection preferentially affects proteins involved in molecular interactions, we collected experimentally supported protein-protein interactions (PPIs) for the investigated proteins from multiple resources and classified their interaction partners as human or non-human (Supplementary Material 10: Table S24). We first examined interactions between human proteins. Interestingly, PSRs showed a depletion in proteins participating in human PPIs; with the direction and magnitude of the effect depending on the background used (Fig. 3/A, Supplementary Material 11: Table S25). Similarly, we observed no consistent enrichment across subcellular localization categories (Supplementary Material 11: Table S26). Connectivity (the number of human interaction partners of a protein) also did not show a simple relationship with positive selection: PSRs were enriched in some proteins with intermediate connectivity but not in proteins within the highest connectivity class (hubs) (Supplementary Material 11: Table S27). Next, we zoomed in on the interactions and examined residues directly involved in experimentally resolved human protein-protein contacts. In contrast to the overall enrichment of PSRs among contacting residues of protein-protein interaction interfaces described in the previous section, PSRs were depleted among residues directly contacting human protein partners in both the HQ_PSR and LS_PSR datasets and relative to both randomized backgrounds (Supplementary Material 11: Table SS28, Fig. 3/A), implying that the previously detected general enrichment in interacting residues (Fig. 2/D,E) might come from inter-species PPI interfaces. Therefore PSRs seem to avoid the residues that maintain direct atomic contacts with human partner proteins, implying that they may modulate human PPIs without abrogating them.

**Figure 3.**
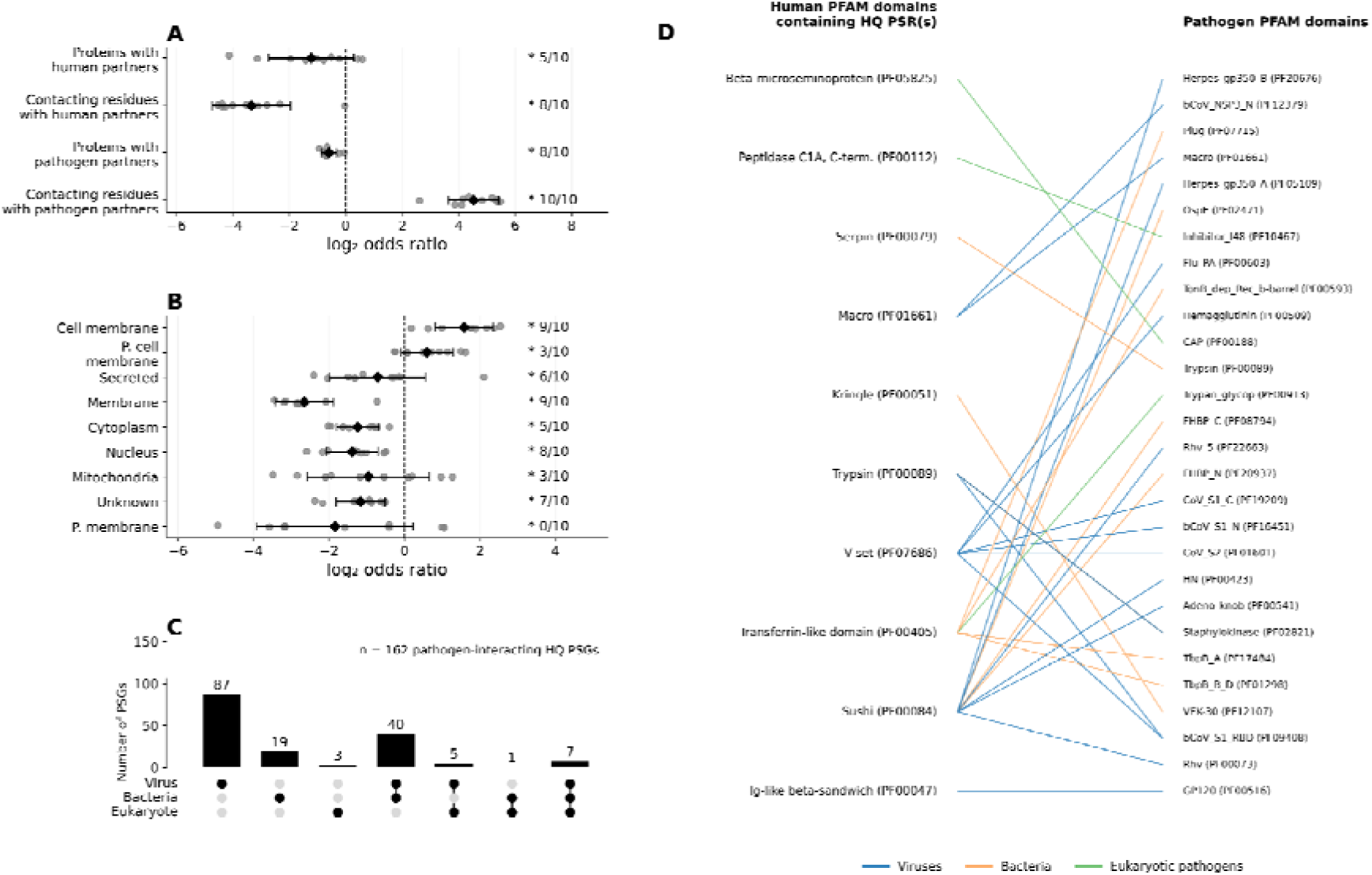
PSRs are enriched in cell-membrane pathogen-interacting proteins and among contacting residues of human-pathogen PPI interfaces. All figures show data from the HQ_PSR dataset. **A:** Forest plots showing the enrichment or depletion of positively selected residues (PSRs) in proteins with human interaction partners, pathogen interaction partners, and in contacting residues of experimentally resolved human and pathogen interaction interfaces compared with the respective random background. **B:** Forest plots showing the enrichment or depletion of PSRs in pathogen-interacting proteins across different subcellular localization categories. For a description of forest plots see Figure 1. **C:** UpSet plot summarizinthe types of pathogen interactions of the 162 positively selected proteins (PSGs) targeted by pathogens. **D:** Pathogen interactions of human domains within PSGs that contain PSRs. Human PFAM domains (left) are connected to pathogen domains or proteins (right), with edge colors indicating pathogen taxonomic groups (blue, viruses; orange, bacteria; green, eukaryotic pathogens).

Despite the lack of enrichment of PSRs in human PPI interfaces, we examined whether a recognizable pattern emerges in how the affected interactions are formed. Although PSRs were not enriched within PFAM domains in general, we asked whether they preferentially occur in domains known to mediate PPIs. No enrichment was observed in PFAM domains with experimentally determined domain-domain interactions nor in domains that were shown to interact in other proteins (Supplementary Material 12: Table S29-30). Because a large fraction of cellular PPIs can be accounted for by short linear motif (SLiM)-mediated interactions [59], which are known to be effectively modulated through evolution [60], we next investigated whether PSRs preferentially localize to known SLiMs or the corresponding SLiM-binding domains. Surprisingly, we observed a depletion of PSRs in these interaction modules (Supplementary Material 13: Table S31), even when predicted SLiMs were also incorporated into the analysis. We further extended the analysis by identifying putative ligand-binding pockets in the AlphaFold-predicted protein models of PSGs using PRANK [61] and observed a depletion of PSRs in them (Supplementary Material 13: Table S31). In all, despite considering domain-domain and ligand-domain interactions (including both the peptide and the domain side), we could not identify a characteristic pattern in the affected human interfaces.

PSRs were also generally depleted in proteins with known pathogen interaction partners (Fig. 3/A, Supplementary Material 14: Table S32) and no relationship could be identified between positive selection and the number of such partners (Supplementary Material 14: Table S32). However, when pathogen interactions were analyzed according to protein localization, PSRs were enriched in pathogen-interacting cell membrane proteins in the HQ_PSR dataset, whereas this enrichment was weaker and less consistent in the LS_PSR dataset (Fig. 3/B, Supplementary Material 14: Table S34). Also, PSRs were enriched in known virus receptor proteins [62] in both datasets (Supplementary Material 14: Table S35), suggesting a role for PSRs in modulating virus-host interactions. Additionally, we observed a strong enrichment of PSRs among residues directly contacting pathogen proteins in experimentally resolved complex structures in both analyzed datasets (Fig. 3/A, Table1, Supplementary Material 14: Table S35). This implies that in contrast to human-human PPIs where, by avoiding directly contacting residues, PSRs seemed to rather play a modulatory role, in the case of certain human-pathogen PPIs, positive selection seems to target the contacting residues and thereby abrogating the interaction.

Finally, we integrated experimentally supported human-pathogen PPIs with PFAM interaction annotations and taxonomic information to identify molecular interfaces potentially targeted by positive selection. Around 50% of PSGs were found to interact with viral, bacterial or eukaryotic pathogens, with some being targeted by multiple pathogen groups (Fig. 3/C, Supplementary Material 15: Table S36). Interestingly, the small set of proteins targeted by all three pathogen groups consists predominantly of extracellular immune regulators and host defense proteins, including CD4, Complement factor H, Plasminogen, Serotransferrin and Alpha-1-antitrypsin (Supplementary Material 15: Table S37). A notable intracellular exception is PPP1R15A, which is known to be hijacked by pathogens [37]. These human proteins participate in fundamental host processes that are recurrently exploited by diverse pathogens. Because interaction databases are biased toward well-studied proteins, this observation should be interpreted with caution. At the domain level, several extracellular interaction modules affected by PSRs repeatedly occurred in pathogen interactions, including Immunoglobulin, Immunoglobulin V-set, Sushi (SCR), Trypsin, and Transferrin domains. These domains were found to interact with proteins from phylogenetically diverse pathogens, including HIV gp120, coronavirus spike proteins, influenza proteins, picornavirus capsid proteins, bacterial transferrin-binding proteins, and staphylokinase family proteins, among others (Fig. 3/D, Supplementary Material 15: Table S38).

**Table 1:** Human-pathogen molecular complexes where human PSRs contact the pathogenic protein.

| Gene name | PSR(s) | PDB(s) | Pathogen group | Pathogen species | Pathogen protein |
|---|---|---|---|---|---|
| PLG | 602, 663 | 1bml | B | Streptococcus dysgalactiae subsp. equisimilis | Streptokinase |
| PLG | 663 | 1bui | B | Staphylococcus phage 42D.m | Staphylokinase |
| CD46 | 103 | 3inb | V | Measles virus strain Edmonston | Hemagglutinin glycoprotein |
| CD46 | 103 | 3l89 | V | Human adenovirus 21 | Fiber protein |
| CD46 | 103 | 8qk3 | V | Human adenovirus 11 | Fiber protein |
| CD4 | 77 | 1g9m, 8fyj… (40 PDB structures) | V | Human immunodeficiency virus 1 | Envelope glycoprotein GP120 |
| CD55 | 170, 178 | 3iyp | V | Echovirus E7 | Capsid protein |
| CD55 | 170 | 6la5 | V | Echovirus E11 | Capsid protein VP2 |
| CFH | 1184 | 5nbq | B | Borrelia burgdorferi | Outer surface protein E |
| CFH | 329 | 4aym | B | Neisseria meningitidis MC58 | Factor H binding protein |
| CTSV | 226 | 3h6s | F | Clitocybe nebularis | Clitocypin analog |
| TF | 382, 625, 564, 576, 591 | 3v89, 3v8x | B | Neisseria meningitidis serogroup B | Transferrin binding protein A (TbpA) |
| CD59 | 67, 73, 85 | 4bik, 5imt | B | Streptococcus intermedius | Intermedilysin |
| CD59 | 37, 67, 73, 85 | 5imy | B | Gardnerella vaginalis | Vaginolysin |
| LTF | 382 | 7jrd | B | Neisseria meningitidis MC58 | Lactoferrin-binding protein B |
| LTF | 382 | 7n88 | B | Neisseria gonorrhoeae | Lactoferrin-binding protein B |

These findings are consistent with the results of previous studies showing that pathogens represent a major selective pressure shaping mammalian and human protein adaptation and genetic variability [27–29], but at the same time provide further insights. Van der Lee *et al.* highlighted some individual examples of positively selected residues located at interfaces with viral proteins without looking for statistical enrichments [23], while Moutinho et al. demonstrated a general enrichment of positively selected residues at protein-protein interaction interfaces using *D. melanogaster* and *A. thaliana* proteins [10], but with no calculations for of host-pathogen interactions. We demonstrated that while human PSRs are depleted in human PPI interfaces, they are significantly enriched among the contacting residues of human-pathogen PPI interfaces across all datasets and randomizations, thereby providing statistically solid proof for human adaptive evolution aiming at neutralizing pathogen attacks in humans.

### Positively selected residues are depleted in pathogenic missense variations but tend to affect known drug targets

To examine the relationship between positive selection and human genetic variation, we collected experimentally verified missense SNVs classified as benign (53 226 variants) or pathogenic (39 821 variants) from UniProt (Supplementary Material 16: Table S39). PSRs showed a lower pathogenic-to-benign mutation ratio, although the residue-level comparisons were only moderately significant in the HQ_PSR dataset because relatively few experimentally annotated variants overlapped PSRs (Supplementary Material 16: Table S40). A stronger and more consistent signal was observed at the protein level: PSGs showed significant enrichment of benign over pathogenic mutations compared with randomly selected proteins (Fig. 4/A, Supplementary Material 16: Table S41).

**Figure 4.**
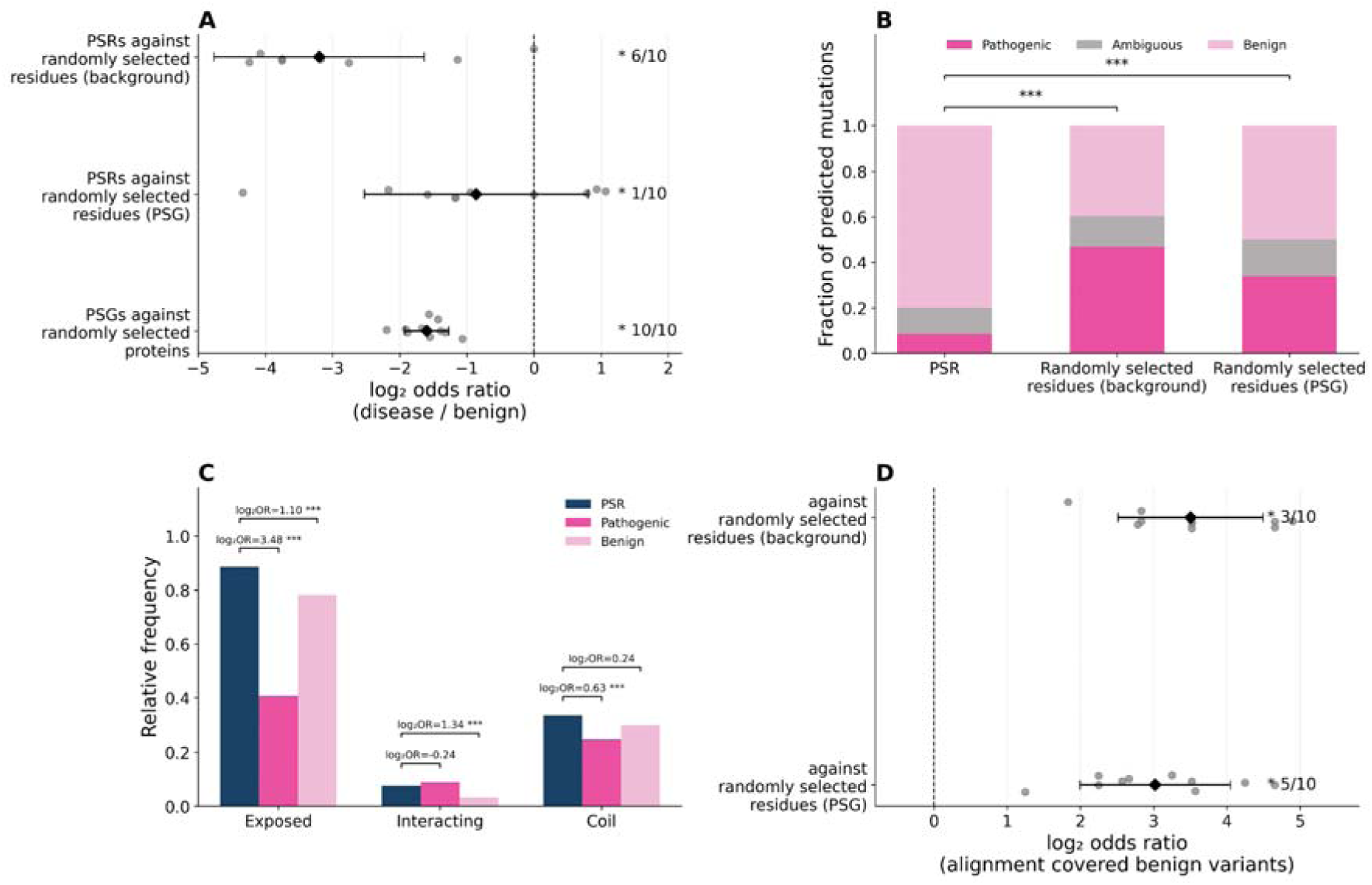
PSRs are depleted in pathogenic missense variants and exhibit distinct structural features from variants. All figures show data from the HQ_PSR dataset. **A:** Forest plot showing the relative depletion of pathogenic missense variants compared with benign missense variants at PSRs and in PSGs. Comparisons include PSRs versus the respective random background, PSRs versus randomly selected residues from PSGs, and PSGs versus randomly selected proteins from the respective background. **B:** Stacked bar plot showing the distribution of AlphaMissense-predicted pathogenic, ambiguous, and benign substitutions at PSRs and the corresponding random background datasets. Values for randomly selected residues represent the mean across randomizations. **C:** Relative frequencies of residues with the indicated structural features among PSRs, experimentally verified pathogenic and benign mutations. **D:** Forest plot showing whether experimentally verified benign mutations at PSRs correspond to amino acid states observed at the same alignment position in homologous PSG sequences. For a description of forest plots see Figure 1. On the bar graphs and violin plots * marks p < 0.05, ** marks p < 0.01 and *** marks p < 0.001 (Pearson’s χ² test).

Because experimentally annotated missense variants provide limited coverage at individual residues, we repeated the analysis using AlphaMissense [63] variant effect predictions. This larger dataset supported the same trend: PSRs were highly significantly enriched in benign predicted mutations and depleted in pathogenic predicted mutations relative to random background residues (Fig. 4/B, Supplementary Material 16: Table S42).

We next compared the structural properties of PSRs to those of experimentally verified pathogenic and benign mutations. Pathogenic mutations were more frequently associated with buried residues, whereas PSRs were highly significantly more exposed than pathogenic mutations in both datasets. The relationship between PSRs and benign mutations was less consistent, with HQ_PSRs showing significantly larger exposure than benign mutations (but LS_PSRs showing lower exposure). PSRs were highly significantly enriched in contacting residues of protein-protein interaction interfaces relative to benign mutations in both datasets, whereas enrichment relative to pathogenic mutations was observed only in the LS_PSR dataset. PSRs were also highly significantly enriched in coil regions relative to pathogenic mutations in both datasets. Thus, PSRs differ consistently from pathogenic variation in structural properties such as exposure and coil localization, while their relationship to benign variation and protein-protein interaction interfaces is dataset dependent (Fig. 4/C, Supplementary Material 16: Table S43).

Finally, we examined experimentally verified benign mutations at PSRs using multiple sequence alignments of PSGs from primates. In the HQ_PSR dataset, the amino acid introduced by the benign mutation was often observed at the equivalent alignment position in homologous sequences, whereas this was less common for randomly selected residues. A similar but weaker and less consistent tendency was observed in the LS_PSR dataset. Although this analysis is limited to a small number of mutations, it suggests that some benign variants at PSRs correspond to amino acid states already sampled during evolution (Fig. 4/D, Supplementary Material 16: Table S44-45).

We also examined whether PSRs affect proteins targeted by drugs that have entered clinical development (based on ChEMBL [51]) (Supplementary Material 17: Table S46). In the larger LS dataset, significantly more PSGs (19.7%, Supplementary Material 17: Table S47) were clinical-stage drug targets than randomly selected proteins. This enrichment was not attributable to the overrepresentation of cell-membrane proteins among PSGs: no enrichment was observed when cell-membrane proteins were considered separately, whereas enrichment was retained among non-cell-membrane PSGs. A similar enrichment was not detected in the smaller HQ dataset. These results suggest an association between positive selection and established pharmacological relevance in the structure-based LS dataset that extends beyond the enrichment of PSGs at the cell membrane.

### Positively selected residues can be distinguished from random residues using sequence and structural features

To evaluate whether the properties identified above are sufficient to distinguish PSRs from non-selected residues, we trained a machine learning model. A Random Forest classifier (Supplementary Material 18: Table S48) was trained on curated features, including protein localization, membrane topology, solvent accessibility, secondary structure, interaction data and mutation annotations.

The HQ_PSR vs HQ_random_background Random Forest models achieved ROC-AUC values of 0.81 and the LS_PSR vs LS_random_background predictors achieved 0.75. The predictive signal was also retained across datasets: models trained on HQ_PSR achieved a ROC-AUC of 0.72 when tested on LS_PSR, while models trained on LS_PSR achieved a ROC-AUC of 0.75 when tested on HQ_PSR (Fig. 5/A, Supplementary Material 18: Table S49).

**Figure 5.**
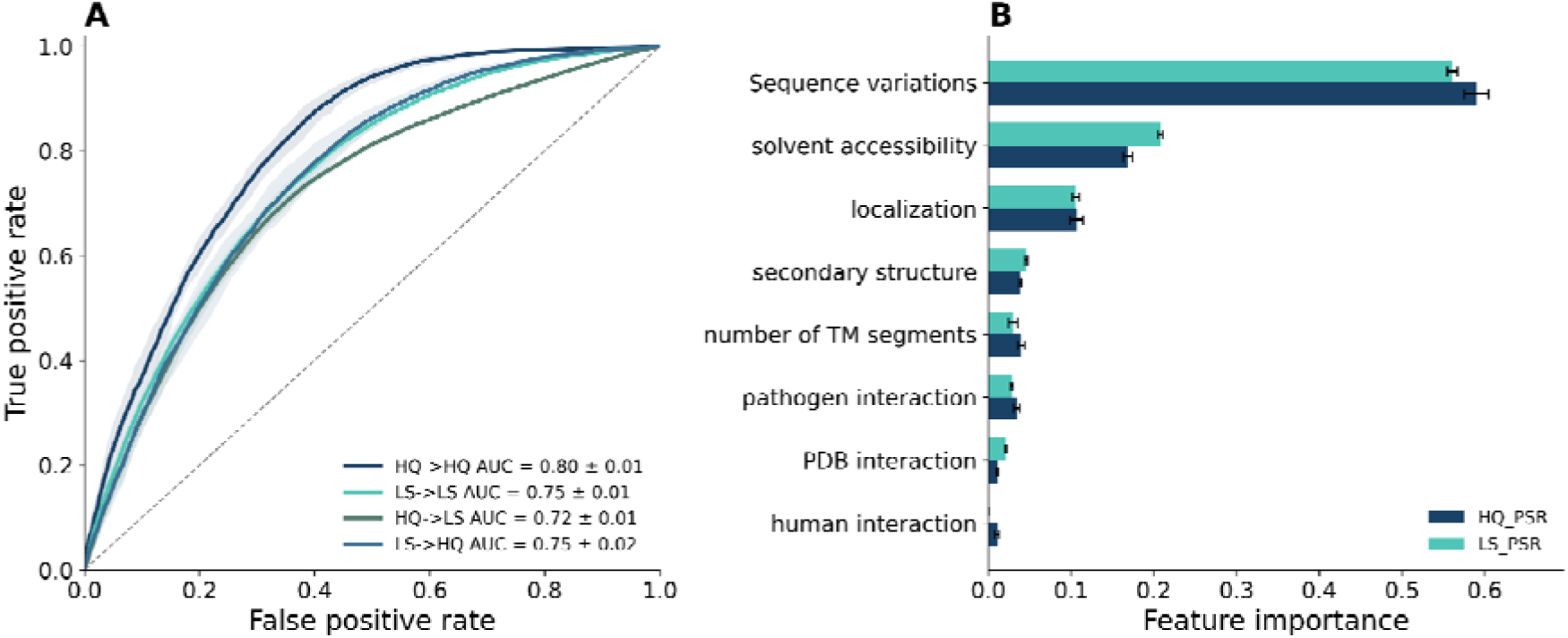
Structural features, sequence variations, localization, and interaction information effectively distinguish PSRs from other residues. A: Performance of a Random Forest classifier trained to distinguish PSRs from randomly sampled residues using structure, localization, interaction, membrane topology, and mutation-related features. The performance metrics of the different models tested on different PSR datasets are indicated. B: Feature importance of the Random Forest model. Feature importance values indicate the contribution of individual feature groups to classification performance.

Considering the HQ_PSR models, feature importance analysis of the Random Forest model revealed that missense variant information contributed most strongly to classification performance. Predicted benign and disease-associated variants accounted for approximately 59% of the total feature importance. Solvent accessibility (16.9%), localization (10.7%), membrane topology (4%) and secondary structure (3.9%) provided additional predictive information, whereas interaction-related features contributed less individually (Fig. 5/B, Supplementary Material 18: Table S50). Similar trends were also visible in the LS_PSR models.

We also performed a feature ablation analysis that confirmed AlphaMissense data to be the strongest predictor (yielded 0.75 and 0.66 AUC on the HQ and LS datasets, respectively), and without AlphaMissense the features retained predictive information (0.63 and 0.62 AUC). Notably, feature importance also heavily depends on the feature type, with continuous residue-level features (e.g., AlphaMissense and accessibility) tending to receive higher importance than binary protein-level ones (e.g., pathogen interactions) (Supplementary Material 18: Table S51).

These results indicate that PSRs possess characteristic sequential and structural properties that distinguish them from random residues. Notably, if the primary goal is to maximize accuracy (rather than demonstrating which features are important), more advanced techniques, such as protein embeddings and neural networks might achieve better performance.

## Discussion

This study provides a comprehensive structural and functional characterization of positively selected residues (PSRs) previously identified in human proteins by two dedicated analyses. We primarily relied on the high-confidence, manually curated human PSR dataset of van der Lee and colleagues [23], but we also used an independent, structure-based, larger PSR dataset published by Slodkowicz and Goldman [25] throughout the analysis. Previous studies typically compared PSRs to all remaining residues or to structural classes within the same proteins. In contrast, we employed two matched randomization schemes: a dataset-dependent wider background based on the sequence pool from which PSRs were originally derived and a narrower background restricted to positively selected genes (PSGs) of the given dataset. This design allowed us to distinguish features associated with positive selection itself from features that merely reflect the properties of proteins containing PSRs. While several of our observations confirm earlier findings, including the preference of PSRs for solvent-exposed regions and their tendency to cluster within protein structures, the substantially expanded structural coverage provided by AlphaFold 2 models allowed us to investigate these properties across a considerably larger set of proteins and thereby gaining more robust and unbiased results. More importantly, combining structural annotations with localization, interaction, and variation data revealed that positive selection is preferentially associated with specific (mainly extracellular) exposed interaction surfaces rather than with particular structural elements or protein domains.

Our results highlight a close relationship between positive selection and proteins involved in extracellular interactions. Accordingly, the analysed PSRs in both datasets were enriched in secreted and cell-membrane proteins, whereas depleted in nuclear proteins. Interestingly, this tendency differs from that reported by Moutinho and colleagues previously, who observed enrichment of nuclear proteins in their combined *Drosophila* and *Arabidopsis* datasets [10]. The opposing trends observed in the extent of adaptive evolution affecting nuclear proteins could be associated with the strong across-species variation in the molecular adaptive rates proposed by many studies [10]. While fruit fly species show substantial rates of adaptive protein evolution [64–67], primates show very low rates [64–68]. This also means that the observed effects and correlations reported in this study cannot be confidently extended to other species with substantially different adaptive rates.

Our localization and interaction analyses consistently indicate that adaptive evolution preferentially targets cell-surface and secreted proteins, which constitute the primary interface between human cells and their environment [23,27]. Consistent with this view, although not enriched in pathogen-interacting human proteins in general, PSRs were significantly enriched at experimentally resolved contacts of pathogen-binding interfaces. Although the relatively small number of available complex structures currently limits this observation, it suggests that adaptive evolution does not primarily target pathogen-interacting proteins *per se*, but rather certain molecular surfaces through which pathogens recognize their hosts are important hotspots for adaptive changes. This interpretation is consistent with the observed enrichment of PSRs at exposed interaction interfaces and with their tendency to form localized surface clusters, which was also shown by Slodkowicz and Goldman [25]. Our findings are also in agreement with earlier studies that identified viruses as an important driver of protein adaptation in mammals [27] and pinpointed pathogens as a major selective pressure shaping human genetic variability and evolution [28,29].

The presence of several recurrent protein domain types in the dataset also supports this interpretation. Immunoglobulin, Sushi repeat and Trypsin domains interact with pathogen proteins and are repeatedly affected by PSRs. In addition, these domains often appear in bitopic membrane proteins (the most affected TMP topology class), including *SLAMF6*, *SLAMF7*, CD33, *CD46*, SIGLEC6, *PIGR, CR2, CD46* and *TMPRSS2* (Fig 6/A). For example, TSPAN8 and EMP1 are TM proteins with 4 membrane spanning regions, with all PSRs confined to extracellular loops (Fig. 6/B). In TSPAN8, the large extracellular EC2 loop is the primary determinant of interaction partner specificity and mediates the assembly of tetraspanin interaction networks [69,70]. Both TSPAN8 and EMP1 possess numerous experimentally verified human interaction partners, suggesting that these extracellular loops constitute major molecular recognition surfaces. Although the interaction mechanism of EMP1 remains less well characterized, it exhibits a similar topology to TSPAN8, with all PSRs localized to extracellular loops.

**Figure 6.**
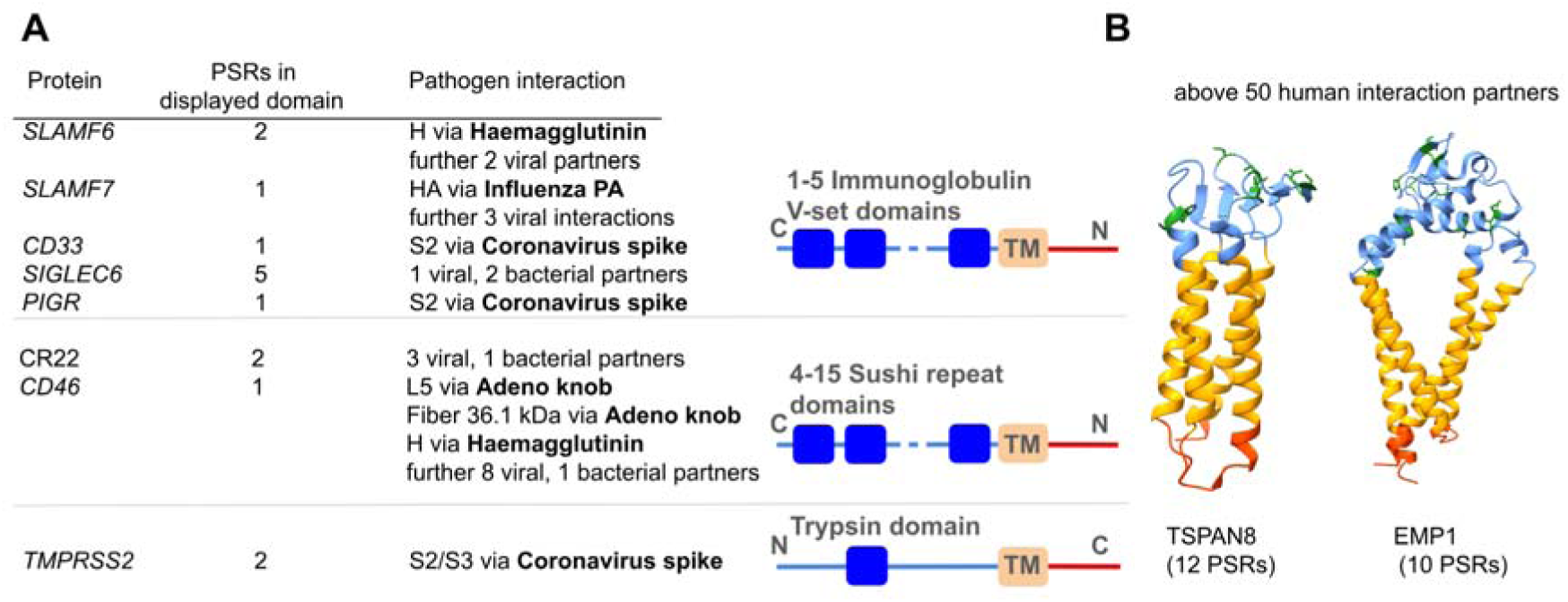
Examples illustrating the structural context and evolutionary consequences of positive selection. A: Representative bitopic transmembrane proteins containing extracellular Immunoglobulin V-set, Sushi repeat, and Trypsin domains that harbor positively selected residues and interact with multiple viral and bacterial pathogen proteins. Several of these extracellular domains represent recurrent targets of pathogen binding. **B:** AlphaFold-predicted structures of TSPAN8 and EMP1 (blue: extracellular, yellow: membrane, red: intracellular, green: PSRs). Membrane definitions were taken from the TmAlphaFold database [75]

An unexpected observation was the absence of enrichment in intrinsically disordered regions (IDRs) and SLiM-mediated interactions. PSRs were claimed to preferentially localize to disordered regions in several species with higher adaptive rates, such as Drosophila [10] and yeasts [17]. Moreover, a previous analysis proposed that human long IDRs are preferential targets of positive selection and estimated a four-fold difference in their relative rate of adaptive substitutions compared to ordered regions [11]. Surprisingly, in the van der Lee dataset we did not observe an enrichment of PSRs within predicted IDRs. This observation does not reflect a structural bias in the dataset, since we tested the pool of more than 11 000 proteins used for PSR detection and it was not biased towards proteins with lower disorder content compared to the whole proteome. Therefore, the disagreement may stem from differences in methodology and alignment depth (primates in the case of Van der Lee *et al.* and mammals in the earlier study by Afanasyeva *et al.*) or other confounding effects. Based on the associated literature, increased surface accessibility and an enrichment in coil structures seem as general traits of PSRs invariably identified across different species and PSR detection approaches, while other features, such as preferential localization to IDRs, showed distinct patterns.

Regarding the typical interaction modules of IDRs, SLiMs, numerous pathogens manipulate host signalling through SLiMs, which can evolve rapidly due to their minimal sequence requirements [71,72]. If rewiring SLiM-mediated interactions represented the dominant mechanism underlying host adaptation to pathogen attacks, one would expect PSRs to accumulate within SLiMs or SLiM-binding domains preferentially targeted by pathogens. Instead, neither feature was enriched in our dataset, and positive selection was primarily associated with PFAM domains shown to interact in other proteins via domain-domain interactions. The lack of PSRs in SLiM-binding domains may reflect that changes within the respective SLiM-binding pockets could not selectively shut off or modulate pathogen attacks without also disrupting the important human PPIs that pathogens hijack through SLiM mimicry [71,72]. Another reason could be the fast pace of co-evolution, meaning that changes within the respective SLiM-binding pockets could easily be compensated by co-evolution of the respective motif on the pathogen’s side and therefore would not provide lasting benefit. Thus, we assume that the host-pathogen domain-domain interfaces delineated by PSRs are more unique to pathogen targeting, making them more ideal for selective changes through adaptive evolution.

Van der Lee *et al.* previously reported that PSGs contain significantly more unique missense variants than expected [23]. By separating benign and pathogenic variants, we show that benign substitutions mainly explain this excess. We observed the same tendency at the level of PSRs on a larger dataset derived from AlphaMissense variant effect predictions: PSRs were enriched in predicted benign and depleted in pathogenic substitutions relative to both applied backgrounds. Benign missense variation also emerged as the most informative feature in the Random Forest classifier built to distinguish PSRs, suggesting that residue-level patterns of human variation capture information associated with positive selection.

The interplay between positive selection, missense sequence variations and pharmacological targeting may have translational implications. PSRs are depleted in pathogenic variants, suggesting that they prefer residues that are more tolerant to amino acid substitutions. This also links to their preference for accessible residues, since those are generally less likely to be pathogenic than buried ones [73]. In parallel, PSGs showed an enrichment among clinical stage drug targets in the LS dataset. This enrichment obviously cannot reflect an effect of drugs on human protein evolution (because 1) the LS dataset was derived from mammalian alignments including only the human reference genome, but no data on human polymorphisms, 2) drugs were developed so recently and are applied so sporadically that they didn’t have enough time to exert a detectable effect on human proteins and 3) only about half of the drug target PSGs have already approved drugs (104 of 206 proteins), the rest is only in the development stage), therefore it rather reflects that adaptive evolution acts on the pool of proteins that can be improved to neutralize certain human diseases, which overlap with the ones selected for drug targeting with a similar purpose. Together, these observations imply that human variations and adaptive changes may provide useful complementary evidence when prioritizing protein regions for therapeutic intervention.

Along these lines, we propose that PSR data could complement current approaches used for the identification of optimal drug targets to neutralize pathogen attacks. To this end, PSRs should be consistently identified in primate and mammalian alignments (at least for the proteins for which one-to-one orthologs can be identified). Human PSGs that are targeted by pathogens (with at least one pathogen PPI) could be selected, their structural complexes obtained or modelled, and their likely pathogen-contacting domain surfaces delineated by PSRs could be searched for druggable pockets [74]. Obviously, surfaces/pockets not known to be involved in essential human interactions and lacking disease-associated mutations should be selected to mitigate potential side effects.

## Conclusions

Overall, our analyses indicate that adaptive evolution in the primate/human lineage preferentially targets extracellular solvent-exposed molecular recognition surfaces. The occurrence of PSRs in folded domains and intrinsically disordered regions is balanced, while among secondary structure types, PSRs are depleted in –helices and enriched in coils. By integrating evolutionary, structural and interaction data, we provide a refined view of the structural determinants of positive selection and identify certain protein families, domain types and interaction interfaces that are recurrently affected by positive selection or serve as hotspots for adaptive mutations and thus represent promising candidates for future functional studies. As the structural coverage of host and host-pathogen molecular interactions and comparative genomic datasets improves, these approaches should support a more comprehensive understanding of how adaptive evolution acts on molecular interactions. At the same time, more precise identification of adaptive substitutions in the future could help delineate human protein surfaces targeted by the subset of human pathogens influential enough to have a detectable impact on human evolution, and thereby may facilitate the mapping of host (and pathogen) protein surfaces eligible for drug targeting. Similarly, as proteins subject to positive selection tend to overlap with clinical stage drug targets, the signal of adaptive changes could be a novel feature to consider when appointing therapeutic targets.

## Data availability

All data are available as supplementary material.

## Additional Files

**Supplementary Material 1:** Supplementary Figures

**Supplementary Material 2:** Positively selected residues and matched randomized residue sets from the van der Lee (HQ_PSR) and Slodkowicz datasets (LS_PSR).

**Supplementary Material 3:** Comparison of protein-protein interaction data, PDB coverage and protein disorder content of the background pool sets and the human proteome.

**Supplementary Material 4:** Subcellular localization and transmembrane topology of positively selected genes.

**Supplementary Material 5:** Subcellular localization of positively selected genes and residues.

**Supplementary Material 6:** Transmembrane topology and membrane localization of positively selected genes and residues.

**Supplementary Material 7:** Association of positively selected residues with protein domains and intrinsically disordered regions.

**Supplementary Material 8:** Structural properties of positively selected residues.

**Supplementary Material 9:** Three-dimensional spatial clustering of positively selected residues.

**Supplementary Material 10:** Protein-protein interaction dataset used in the interaction analyses.

**Supplementary Material 11:** Association of positively selected residues with human protein-protein interactions.

**Supplementary Material 12:** Association of positively selected residues with interacting protein domains.

**Supplementary Material 13:** Association of positively selected residues with short linear motifs and other functional sequence features.

**Supplementary Material 14:** Association of positively selected residues with pathogen protein-protein interactions.

**Supplementary Material 15:** Pathogen interactions and domain-level interaction features of positively selected genes and residues.

**Supplementary Material 16:** Relationship between positive selection and human missense variation.

**Supplementary Material 17:** Drug-target potential of positively selected genes.

**Supplementary Material 18:** Random-forest prediction of positively selected residues from protein features.

## Author contributions

R.P. and L.D. designed the study. L.D., E.S, E.F, A.S., A.T. and R.P. performed all analyses. L.D. and R.P drafted the manuscript. L.D, A.T., G.E.T. and R.P. reviewed and edited the manuscript. R.P, L.D, A.T. and G.E.T. obtained funding for the project. All authors read and approved the final manuscript, all authors reviewed the manuscript.

## List of abbreviations

IDR: Intrinsically Disordered Region
HQ: High Quality
LS: Large Scale
P. Membrane: Peripheral Membrane
PSR: Positively Selected Residue
PSG: Positively Selected Genes
PDB: Protein Data Bank
PPI: Protein-protein interaction
SLiM: Short linear motif
SNV: Single nucleotide variation
TM: Transmembrane
TMP: Transmembrane Protein

## Acknowledgement

We thank Greg Slodkowicz and Nick Goldman for sharing the dataset used in their study. We also thank Robin van der Lee for sharing details and data of their earlier study.

## Ethics declarations

### Ethical approval

Not applicable.

### Consent to participate

Not applicable.

### Consent for publication

Not applicable.

