## Supplementary Material 1 for "Positive selection tends to act on exposed extracellular regions and delineate interaction modules targeted by pathogens"

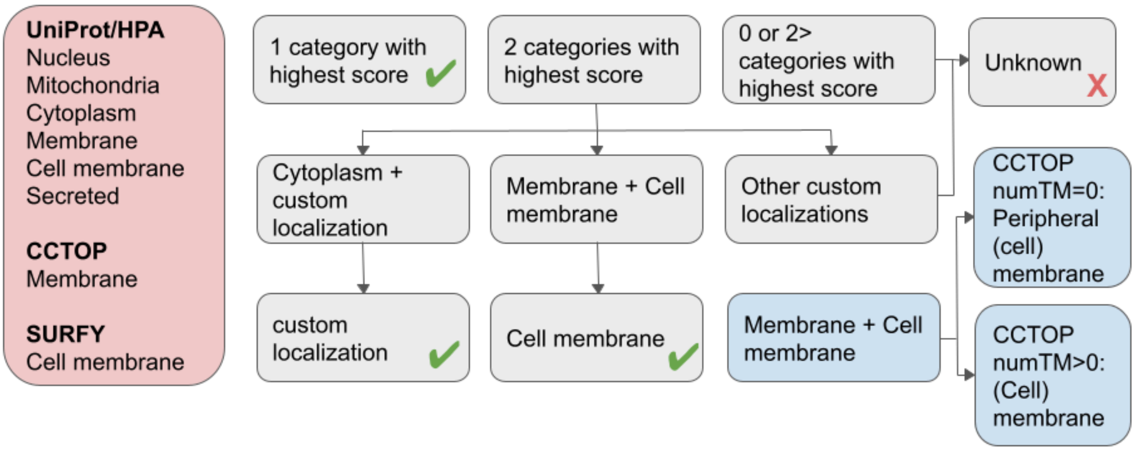


**Figure S1: Schematic figure of how localizations were assigned to proteins.** The data sources used for localization annotation are shown on the left (light red box), along with the localization categories supported by each source. The localization assignment procedure is illustrated by grey boxes. For proteins assigned to the membrane or cell membrane categories, CCTOP topology predictions were used to distinguish transmembrane from peripheral proteins (blue boxes).

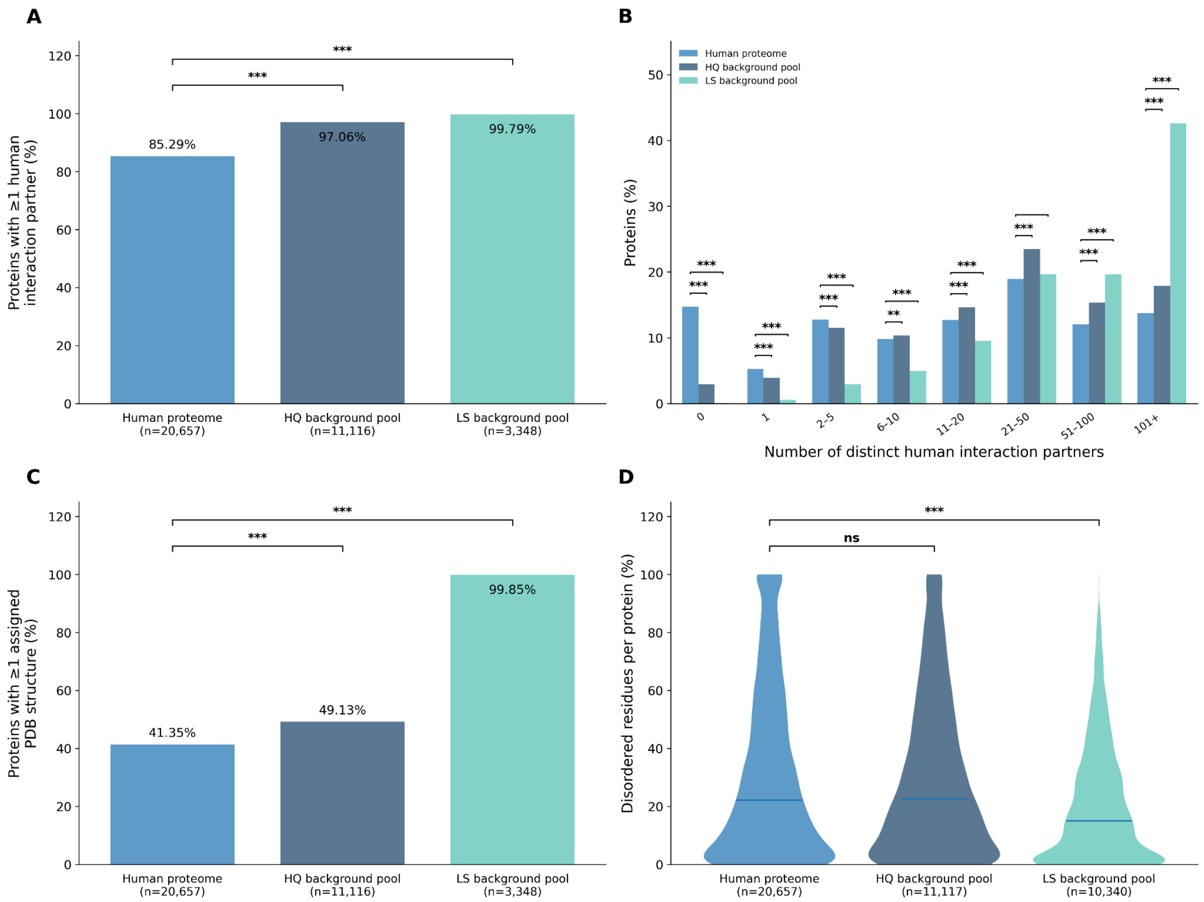


**Figure S2: Protein-interaction, structural coverage and disorder content characteristics in the background pool sets and in the human proteome.** A: Number of proteins with at least one human interaction partner in the background pool sets and in the human proteome. B: Human interaction partners/proteins in the background pool sets and in the human proteome. C: Proteins with at least assigned PDB structures in the background pool sets and in the human proteome. D: Protein disorder content in the background pool sets and in the human proteome. On the bar graphs (Pearson's $\chi^2$ test) and violin plots (two-sided Mann-Whitney tests) * marks $p < 0.05$, ** marks $p < 0.01$ and *** marks $p < 0.001$.

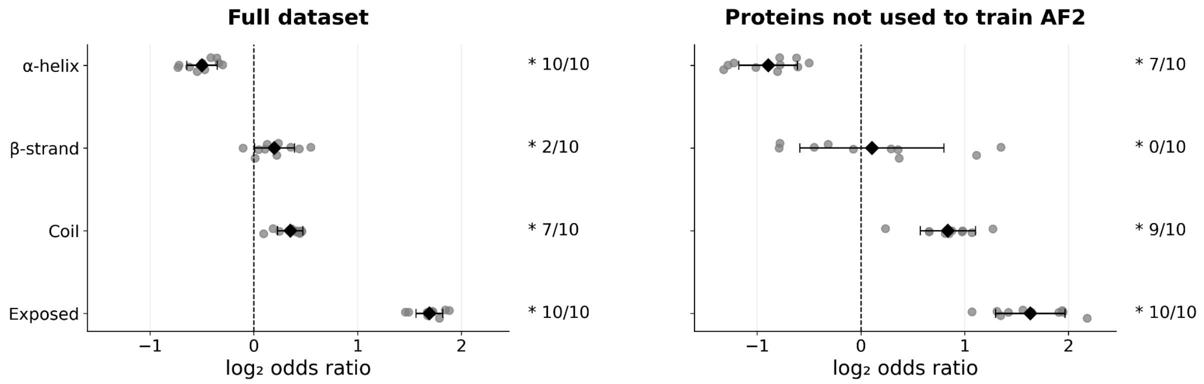


**Figure S3: Structural characteristics of PSRs in AlphaFold2 structures.** Forest plots showing the enrichment or depletion of secondary structure and exposed residues in all AlphaFold2 structures (left) and in those that were not part of the training set (right). On the forest plots, individual dots represent enrichment estimates obtained from 10 independent randomization experiments. Diamonds indicate the mean $\log_2$ odds ratio (expected effect size), while error bars denote the standard deviation across randomizations. On the forest plots the accompanying text reports the fraction of randomizations yielding significant enrichment or depletion at $p < 0.05$ (*: significant) using Pearson's $\chi^2$ test.
